# Investigating impacts of marine sponge derived mycothiazole and its acetylated derivative on mitochondrial function and aging

**DOI:** 10.1101/2023.11.27.568896

**Authors:** Naibedya Dutta, Joe A Gerke, Sofia F Odron, Joseph D Morris, Adam Hruby, Toni Castro Torres, Sarah J Shemtov, Jacqueline G Clarke, Michelle C Chang, Hooriya Shaghasi, Marissa N. Ray, Maxim Averbukh, Sally Hoang, Maria Oorloff, Athena Alcala, Matthew Vega, Hemal H Mehta, Max A Thorwald, Phillip Crews, Marc Vermulst, Gilberto Garcia, Tyler A Johnson, Ryo Higuchi-Sanabria

**Affiliations:** Leonard Davis School of Gerontology, University of Southern California, Los Angeles, CA 90089, United States; Department of Natural Sciences & Mathematics, Dominican University of California, San Rafael, CA 94901, United States; Department of Chemistry & Biochemistry, University of California, Santa Cruz, Santa Cruz, CA, 95064, United States

**Keywords:** mitochondria, aging, stress, C. elegans, cancer

## Abstract

Small molecule inhibitors of the mitochondrial electron transport chain (ETC) hold significant promise to provide valuable insights to the field of mitochondrial research and aging biology. In this study, we investigated two molecules: mycothiazole (MTZ) - from the marine sponge *C. mycofijiensis* and its more stable semisynthetic analog 8-*O*-acetylmycothiazole (8-OAc) as potent and selective chemical probes based on their high efficiency to inhibit ETC complex I function. Similar to rotenone (Rote), a widely used ETC complex I inhibitor, these two molecules showed cytotoxicity to cancer cells but strikingly demonstrate a lack of toxicity to non-cancer cells, a highly beneficial feature in the development of anti-cancer therapeutics. Furthermore, in vivo experiments with these small molecules utilizing *C.elegans* model demonstrate their unexplored potential to investigate aging studies. We observed that both molecules have the ability to induce a mitochondria-specific unfolded protein response (UPR^MT^) pathway, that extends lifespan of worms when applied in their adult stage. Interestingly, we also found that these two molecules employ different pathways to extend lifespan in worms. Whereas MTZ utilize the transcription factors ATFS-1 and HSF-1, which are involved in the UPR^MT^ and heat shock response (HSR) pathways respectively, 8-OAc only required HSF-1 and not ATFS-1 to mediate its effects. This observation underscores the value of applying stable, potent, and selective next generation chemical probes to elucidate an important insight into the functional roles of various protein subunits of ETC complexes and their regulatory mechanisms associated with aging.

## INTRODUCTION

Mitochondria are double membrane bound cellular organelles popularly recognized as the “powerhouse of the cell” for its most well-known function of cellular respiration. Respiration involves a series of protein complexes that transfer electrons from metabolic end products such as NADH and FADH_2_ to a final electron accepter oxygen. These redox reactions of electron transfer are coupled with the establishment of an electrochemical gradient across the inner membrane of mitochondria, which leads to the production of energy in the form of ATP (Casanova et al. 2023). Apart from energy production, mitochondria are also involved in numerous biochemical events in a cell, including signal transduction, metabolism and nutrient breakdown, stress response, immune response, regulation of reactive oxygen species, and calcium homeostasis (Shen et al. 2022; Y. Chen, Zhou, and Min 2018). Additionally, they also play crucial roles in the regulation of cellular fate, including the control of different cell death mechanisms, such as apoptosis (Green 2022). Moreover, maintaining mitochondrial homeostasis is extremely important to achieve optimal cellular function. This homeostatic condition is susceptible to the exposure of various intracellular or extracellular stress conditions, such as oxidative damage, protein misfolding, and mutations in mitochondrial DNA.

When exposed to stressors, cells activate a specific and complex mitochondrial stress response to maintain the integrity and restore the functionality of mitochondria. The key feature of the mitochondrial stress response is the activation of the mitochondrial unfolded protein response (UPR^MT^) that includes the upregulation of genes, which encode essential proteins to restore mitochondrial homeostasis (Bar-Ziv, Bolas, and Dillin 2020). These include genes encoding mitochondria-specific chaperones and proteases to refold or clear damaged mitochondrial proteins. A growing number of studies in the last three decades have established that the combination of mitochondrial dysfunction and an ineffective mitochondrial stress response is strongly associated with the progression of aging and the onset of age-related degenerative diseases like neurodegenerative disorders, cardiovascular diseases, and metabolic disorders. These critical functions of mitochondria mark mitochondrial dysfunction as one of the 12 hallmarks of aging (López-Otín et al. 2023; Amorim et al. 2022; Srivastava 2017).

Despite the consequences of mitochondrial dysfunction, exposure to small doses of mitochondrial stress can be beneficial to an organism. While excessive or chronic mitochondrial stress leads to detrimental consequences to the cell, exposure to lower levels of stress, which can be induced by various interventions such as exercise, dietary restriction, or pharmacological agents, can actually trigger a hormetic response in mitochondria (i.e., mitohormesis) (Ristow and Zarse 2010). Mitohormesis involves the activation of beneficial transcriptional programs, including the UPR^MT^, which improves cellular resilience to subsequent exposure to stress. Notably, several studies have also demonstrated that mitohormesis can even increase lifespan in various model organisms including *C. elegans* and *D. melanogaster* (Dutta, Garcia, and Higuchi-Sanabria 2022; Owusu-Ansah, Song, and Perrimon 2013).

Altogether, understanding mitohormesis can provide a detailed insight into how the cellular stress response pathways can be employed to promote healthspan and increase longevity. However, most models of mitohormesis including the nematode model, *C. elegans*, involve genetic perturbations of mitochondrial function (Higuchi-Sanabria et al. 2018; Durieux, Wolff, and Dillin 2011). While these studies show robust activation of UPR^MT^ and significantly increased longevity upon disruption of mitochondrial function, genetic manipulation may be more limiting in higher organisms, especially in humans. As an alternative to genetic approaches, selected small molecule inhibitors isolated from natural sources have garnered considerable attention due to ease of delivery and their robust downstream effects (Bonkowski and Sinclair 2016). The use of small molecule chemical probes to accelerate the understanding of mitochondrial dynamics has been substantial (C. Ma, Xia, and Kelley 2020). Surprisingly, a number of non-specific chemical probes have and continue to gain widespread use throughout the field of chemical biology (Arrowsmith et al. 2015). Alternatively, a small but growing number of venerable compounds exist that are now proven to be stable (>6 mo. shelf life), potent (<100 nM), and selective for their mechanism of action(Antolin, Workman, and Al-Lazikani 2021). Marine natural products are a burgeoning source of next generation chemical probes to accelerate the study of the cytoskeleton, neuroscience (Johnson et al. 2017), and mitochondrial dynamics (Morgan et al. 2010) in biomedical research.

In this study, we sought to investigate the rarely studied, but potent marine sponge-derived natural product mycothiazole (MTZ), and its semisynthetic derivative 8-*O*-acetylmycothiazole (8-OAc) for their ability to serve as distinct chemical probes that affect the aging process. Recently MTZ was reported as a potent cytotoxic compound against a wide range of cancer cells (Johnson et al. 2020). Both MTZ and 8-OAc were also reported to function as potent and selective inhibitors of the mitochondrial electron transport chain (ETC) complex I (Morgan et al. 2010).

Similar to genetic interventions, we found that MTZ and 8-OAc were able to achieve a mitohormetic effect in *C. elegans* and prolong longevity in adult worms. Perhaps most surprising, we found that these compounds utilized distinct mechanisms to prolong longevity, suggesting that even compounds of the same chemotypes that have identical targets may activate different pathways as chemical probes based on subtle changes in their structural framework. In addition, we found that both compounds could preferentially target cancer cells for the initiation of cell death pathways, while having minimal effects on viability of non-cancer cells. Here, our results not only shed light on how these next generation small molecule chemical probes impact longevity in the model organism, *C. elegans,* but also provide an insight into their mode of actions in human cells.

## MATERIALS AND METHODS

### Human cell lines and maintenance

Human hepatocellular carcinoma (Huh7) and karyotypically normal human fibroblast (BJ) cells were kind gift from Prof. Andrew Dillin’s lab at UC Berkley. Human embryonic kidney 293 (HEK293) cells are a kind gift from Dr. Marc Vermulst’s lab at USC. Cells were cultured in high glucose DMEM media (Cytiva) supplemented with 10% FBS (Cytiva), 1X Penicillin/Streptomycin (VWR Life Science), 1X non-essential amino acid (Cytiva) at 37°C along with 5% CO_2_ in a humified incubator (Thermo). Cells were treated with compounds at ∼70% confluency.

### Cytotoxicity assay

Cytotoxicity of cells was measured using 3-(4,5-diethylthiazol-2-yl)-2,5-diphenyltetrazolium bromide (MTT) assay. 70% confluent were cells grown in multi-well plates and treated with compounds or vehicle/DMSO for 24 h under above mentioned standard condition. Post-treatment growth media was replaced with 0.5 mg/ml MTT solution and incubated for 4h at 37 °C. After removing the MTT solution formazan crystals were dissolved in DMSO and OD was measured at 570 nm using SpectraMax M2 microplate reader.

### Apoptosis assay

Cells were grown up to ∼70% confluency in a 6 well plate and treated with indicated concentration of compounds or vehicle for 24 h. Post incubation cells were processed for apoptosis using flow cytometry according to the protocol described in FITC Annexin-V apoptosis detection kit (BD Pharmingen).

### Measurement of ROS by DHE staining

After treating the cells with indicated concentration of compounds or vehicle control for 24 h growth media was replaced with 5 mM DHE solution and further incubated for 30 mins at 37 °C. ROS amount was determined by measuring the fluorescence of 2-hydroxyethidium, an oxidized product of DHE using flow cytometry.

### Mitochondrial labeling using Mitotracker green dye

70% confluent Huh7 and BJ fibroblast cells, grown on 35mm glass bottom plates were treated with MTZ, 8-OAc or Rote for 24 h. Post treatment growth media was replaced with new media supplemented with 125nM Mitotracker green solution (Thermo) and DAPI and incubated them for 30 mins at 37°C. After incubation cells were imaged using Stellaris 5 confocal microscope.

### Measurement of oxygen consumption rate

Real time oxygen consumption rate was measured by using XF24/96 Extracellular flux analyzer (seahorse bioscience). For this assay, XF assay specific 96 well tissue culture plates (Provided by Agilent) were used to grow Huh7 and BJ cells. First, equal number of cells were grown directly in XF 96 well tissue culture plates and at 70% confluency were treated with 10μM of MTZ, 8-OAc or Rote for 24 h. Post treatment cells were first washed one time with prewarmed XF real-time ATP rate assay media and replaced with 200 µl fresh media. Next the plate was placed in a 37 °C incubator not supplemented with CO_2_ for 30 min. After incubation, media was removed and replaced with fresh prewarmed XF assay media to a final volume 180 µl and measurements were taken. Maximum mitochondrial respiratory capacity was estimated by challenging the cells with 10 µM Oligomycin, 10 µM FCCP, or 5 µM Rotenone. Data was analyzed by seahorse Wave desktop software provided by Agilent.

### Human cell RNAseq analysis

Huh7 and BJ fibroblast cells, cultured to 70% confluency in 6-well plates, underwent treatment with the indicated concentration of compounds or vehicle control and incubated for 24 h. Post incubation growth medium was removed, and cells were collected in 1X PBS in 1.5ml centrifuge tubes. After a PBS wash 500 µl TRIzol was added to the cells followed by the addition of 300 µl chloroform in a heave gel phase-lock tube (VWR, 10847-802) for phase separation using centrifugation. The isolated aqueous phase, enriched with RNA, underwent purification using a standard RNA purification kit. (Quantabio, Extracta Plus 95214-050) as per manufacturer’s directions.

Library preparation (mRNA library, poly A enrichment) and RNA sequencing (NovoSeq PE150. 6G raw data per sample) was performed at Novogene. Three biological replicates were measured per condition.Reads were trimmed with trim_galore-0.6.5-1 and mapped to Genome assembly: GRCh38.p14 with STAR-2.7.3a (Dobin et al. 2013). Mapped reads were counted to genes using feature Counts (Subread-2.0.0) (Liao, Smyth, and Shi 2019) and Homo_sapiens.GRCh38.108.gtf gene annotation. Unwanted variations were removed with RUVSeq-1.32.0 (Risso et al. 2014) and differential expression analysis was performed with DESeq2-1.38.3 (Love, Huber, and Anders 2014) using R-4.2.2.2.

### C. elegans strains and maintenance

All strains used in this study are derivates of the N2 wild-type worm from the Caenorhabditis Genetics Center (CGC) and are listed in **Tab. s3**. For maintenance of worms, animals are grown at 15 °C on OP50 *E. coli* B strain. For all experimental purposes, animals are grown at 20 °C on HT115 *E. coli* K strain bacteria unless otherwise noted. HT115 bacteria carry a pL4440 empty vector (used as a control) or pL4440 vector carrying a partial gene sequence against a specific target gene for RNAi purposes. All experiments are performed on age-matched animals synchronized using a standard bleaching protocol as previously described (Bar-Ziv et al. 2020). Briefly, animals are collected from plates using M9 solution (22 mM KH_2_PO_4_ monobasic, 42.3 mM Na_2_HPO_4_, 85.6mM NaCl, 1 mM MgSO_4_) and bleached using a 1.8% sodium hypochlorite and 0.375 M KOH solution. Intact eggs were washed 4x with M9 solution and L1 synchronization was performed by floating eggs in M9 solution at 20 °C overnight. Synchronized L1s were grown on RNAi plates (1 mM CaCl_2_, 5 µg/mL cholesterol, 25 mM KPO_4_, 1 mM MgSO_4_, 2% agar w/v, 0.25% Bacto-Peptone w/v, 51.3 mM NaCl, 1 µM IPTG, and 100 µg/mL carbenicillin; HT115 *E. coli* K strain containing pL4440 vector control or pL4440 with RNAi of interest). All aging experiments were performed on plates supplemented with 100 µL of 10 mg/mL FUDR spotted directly on the bacterial lawn.

For drug treatments, plates were supplemented with 1, 3, or 5 µM MTZ, 8-OAc and Rote, or an equivalent volume of DMSO. For 1 and 3 µM drug conditions, animals were placed on drug plates from L1. For 5 µM drug conditions, animals were moved onto drug plates from day 1 of adulthood; for all RNAi conditions on 5 µM drugs, L1 animals were grown on standard RNAi plates and moved onto RNAi plates containing drugs at day 1 of adulthood. For drug titration experiments, all animals were grown on all concentrations of drugs from the L1 stage and imaged when control animals reached day 1 of adulthood (3 days at 20 °C).

### C. elegans microscopy

For all transcriptional reporter imaging (*hsp-6p::GFP, hsp-4p::GFP*, *hsp-16.2p::GFP*), synchronized animals were imaged at day 1 of adulthood. Animals were picked off plates and onto standard NGM plates without bacteria containing 5 µL of 100 mM sodium azide to paralyze worms. Paralyzed worms were lined up with the pharynx facing up and imaged on a Leica M205FCA automated fluorescent stereomicroscope equipped with a standard GFP filter and Leica K5 camera and run on LAS X software. Three biological replicates were performed per experiment and one representative replicate image is shown in each figure. Detailed protocols are available at (Bar-Ziv et al. 2020).

### C. elegans RNAseq analysis

For RNA-seq experiments, germline less *glp-4(bn2)* animals were used to avoid progeny contamination and germline effects. Synchronized animals were grown on HT115 bacteria from hatch on standard RNAi plates at 22 °C until day 1 of adulthood. We opted to use 22 °C instead of the standard restrictive 25 °C for *glp-4(bn2)* animals to minimize heat-stress associated with growth at 25 °C. Importantly, after backcrossing these animals 6x to our N2 line, we found that our backcrossed *glp-4(bn2)* animals were more sensitive to elevated temperatures and exhibited full sterility at 22 °C. Animals were collected in M9 solution, M9 was replaced with 1 mL of Trizol and sample was flash-frozen in liquid nitrogen. To isolate RNA, animals were freeze/thawed 3x with liquid nitrogen with a 30 sec vortexing step between each freeze to lyse all animals. 300 µL of chloroform was added to the sample and aqueous separation of RNA was performed using centrifugation in a heave gel phase-lock tube (VWR, 10847-802). The aqueous phase was then applied to a standard RNA purification kit (Quantabio, Extracta Plus 95214-050) as per manufacturer’s directions.

Library preparation (mRNA library, poly A enrichment) and RNA sequencing (NovoSeq PE150. 6G raw data per sample) was performed at Novogene. Three biological replicates were measured per condition. Reads were trimmed with trim_galore-0.6.5-1 and mapped to WBcel235 with STAR-2.7.3a (Dobin et al. 2013). Mapped reads were counted to genes using feature Counts (Subread-2.0.0) (Liao, Smyth, and Shi 2019) and Caenorhabditis_elegans.WBcel235.107.gtf gene annotation. Unwanted variations were removed with RUVSeq-1.32.0 (Risso et al. 2014) and differential expression analysis was performed with DESeq2-1.38.3 using R-4.2.2.2 (Love, Huber, and Anders 2014).

### C. elegans lifespan

All lifespan assays were performed on standard RNAi plates with HT115 bacteria at 20 °C as previously described (Castro Torres et al. 2022). For drug treatments, animals were grown on RNAi plates containing 1 or 3 µM mycothiazole, 8-O-acteyl mycothiazole, rotenone, or an equivalent volume of DMSO from the L1 stage; or animals were grown on standard RNAi plates without drug from the L1 stage and moved onto plates containing 5 µM mycothiazole, 8-O-acteyl mycothiazole, rotenone, or an equivalent volume of DMSO at the day 1 adult stage. All animals were exposed to FUDR from the day 1 adult stage to remove eliminate progeny. The viability of animals were quantified every other day until all animals were scored as either dead or censored. Censored animals are defined as those that exhibit intestinal leakage out of the vulva, bagging (vivipary), desiccation on the walls of the petri dish, or other age-unrelated deaths. Survival curves were plotted using Prism7 software and LogRank statistical testing for calculation of p-values and a minimum of 3 biological replicates were performed for each experiment with all raw data available in **Tab. s4**.

### C. elegans seahorse assay

Synchronized animals were collected from plates using M9 and animals were poured onto an empty NGM plate (no bacteria). ∼10-15 adults worms were pipetted off of these plates while avoiding all progeny and pipetted into each well of a Seahorse XF96 cell culture microplate. Basal oxygen consumption rate was measured using a XFe96 sensor cartridge on a Seahorse XFE96 Analyzer with 2 minutes mixing, 30 second wait, and 2 minutes measuring. Oxygen consumption rate was normalized for number of worms. For low concentration MTZ, 8-OAc, and Rote (1 and 3 µM), animals were grown on plates containing DMSO or compounds from L1 and oxygen consumption rate was measured at day 1 of adulthood. For high concentration (5 µM), animals were grown on standard NGM plates from L1 and moved onto DMSO, MTZ, 8-OAc, or Rote containing plates at day 1 of adulthood for 24 hours, and oxygen consumption rate was measured at day 2 of adulthood.

### Chemical isolation of Mycothiazole (MTZ) and 8-O-acetylmycothiazole (8-OAc)

Biological material collection and identification: Specimens of the marine sponge *C. mycofijiensis* were collected via scuba diving in Vanuatu, as previously documented (Sonnenschein et al. 2006; Morgan et al. 2010). Taxonomic identification was conducted by comparing characteristic biological features with reference samples from the UC Santa Cruz sponge repository. Voucher specimens and underwater photographs can be provided upon request.

Extracts of *C. mycofijiensis* were processed according to previously reported methods (Takahashi-Ruiz et al. 2022; Johnson et al. 2020; Morris et al. 2022). While the fat hexanes (FH) extracts are not commonly processed due to their high lipophilic content, this extract served as an enriched source of mycothiazole (MTZ) with minor amounts of the latrunculin or fijianolide (alternatively referred to as laulimalide) chemotypes. The approximate percentage of mycothiazole in the FH extract was 4.5%. The fat dichloromethane (FD) extract and dichloromethane methanol fat (DMMF) extract were also used for scale up purification and contained approximately 12% and 10% MTZ by weight, respectively. All three extracts (FH, FD and DMMF) were used in the repeated scale-up HPLC isolation of pure MTZ (79.4 mg), for semi-synthesis (54.5 mg) and further biological evaluation experiments (24.9 mg).

HPLC purification was performed on a semi-preparative column (Phenomenex Inc. Luna© 5µm C18(2) 100 Å 10 × 250 mm) in conjunction with a 4.0 × 3.0 mm C18 (octadecyl) guard column and cartridge (holder part number: KJ0-4282, cartridge part number: AJ0-4287, Phenomenex Inc., Torrance, CA, USA). A reversed-phased linear gradient was employed (30:70 CH3CN/H2O to 80:20 over 50 minutes, ramping up to 100% CH3CN from 51 to 61 minutes, then returning to 30:70 for re-equilibration from minutes 62 to 73). Compound detection was measured with an Applied Biosystems 759a UV detector at a single wavelength λmax = 230 nm. For each of the extracts, MTZ would begin to elute at approximately 46 minutes. All purified compounds were dried under an N2 stream, stored in amber vials, and purged with gaseous argon then sealed in a dark desiccator under vacuum.

Pure 8-*O*-acetylmycothiazole (8-OAc) was obtained by two distinct methods. The first method of acetylation employed 29.1 mg of the FH crude extract oil immersed in 1000 μL of acetic anhydride and 1000 μL of dry pyridine that were added to a reaction vial. The mixture was stirred for 24 hours before the reaction was quenched with 1000 μL of di-H2O and 1000 μL of dichloromethane (DCM). The layers were inverted and allowed to separate before the DCM layer was removed from the reaction vial and dried under an N2 stream. The dried DCM layer was purified using RP-HPLC (Phenomenex Luna© 5µm C18(2) 100 Å 10 × 250 mm) to yield one major fraction determined to be 9.9 mg of 8-OAc, resulting in a 34% yield. The second method of acetylation followed the same procedures but rather than reacting with the FH crude extract, purified MTZ (>95%, HPLC) was used. This reaction with 25.4 mg of pure MTZ generated 10.94 mg of the 8-OAc with a 43% yield. Notably, these acetylation reactions demonstrated higher efficiency when utilizing pure MTZ sourced from the FD crude extract. Subsequently, the purity of all compounds was confirmed to be >95% by 1H NMR analysis using a Bruker instrument, featuring a 5 mm triple-resonance cryoprobe (^1^H, ^13^C, ^15^N) operating at 600 MHz for ^1^H experiments (**Fig. s1**). In total, 24.9 mg of MTZ and 20.8 mg of 8-OAc were used in the aforementioned biological evaluation experiments.

### Statistical analysis

For the cell culture experiments all the individual data points combined with mean ± SD and graphs were plotted and statistically analyzed by One-way ANOVA, using GraphPad Prism 10.0. For the nematode lifespan experiments Kaplan-Meier survival graphs were plotted and the Logrank (Mantel-Cox) tests were performed using GraphPad Prism 10.0. ns = not significant, * = p < 0.03; ** = p <0.002; *** = p<0.0002; **** = p< 0.0001

## RESULTS

### Mycothiazole and its acetylated analog show selective toxicity towards cancer cells while exhibiting mitochondrial complex I inhibition for both cancer and non-cancer cells

Utilizing MTZ and its more stable acetylated derivative 8-OAc (Johnson et al. 2020), we aimed to determine the effectiveness of these two compounds against human hepatocellular carcinoma cells (Huh7) and compared these results with two human non-cancer cell types – skin fibroblast (BJ) and kidney epithelial cells (HEK293). Here, we purified MTZ from the marine sponge *C. mycofijiensis* using HPLC purification as previously reported (Johnson et al. 2020; Sonnenschein et al. 2006; Morgan et al. 2010; Morris et al. 2022; Takahashi-Ruiz et al. 2022). 8-OAc was obtained by immersing fat hexane extracts in acetic anhydride and dry pyrimidine for 24 hours before the reaction was quenched with dichloromethane. Purity of all compounds used in this study were confirmed to be >95% by NMR analysis (**Fig. s1**).

In addition, we compared our results to rotenone (Rote), a classically utilized mitochondrial complex I inhibitor chemical probe as a comparative reference.

To study the biological activity of MTZ, 8-OAc, and Rote, we first determined the half-maximal inhibitory concentration (IC50) values for each compound using an MTT assay, which indirectly measures cell viability through a colorimetric assay for cell metabolic activity (Dutta et al. 2022). While all three compounds display high cytotoxicity towards cancer cells, 8-OAc displays much lower cytotoxicity towards non-cancer cells compared to Rote and MTZ (**Fig. 1A, Fig. s2A**). Since mitochondrial dysfunction is often corelated with the production of reactive oxygen species, which can lead to the initiation of apoptosis, we performed an apoptosis assay through annexin-V-FITC, and PI staining. Consistent with MTT assays, all three compounds robustly induced apoptosis in cancer cells. Although Rote showed the highest induction of apoptosis in cancer cells, Rote also induced a noticeable level of apoptosis in non-cancer cells (40-50% at 50 µM), whereas MTZ and 8-OAc failed to induce apoptosis at similar concentrations (**Fig. 1B**). We then directly measure the ROS production capacity of all three compounds by performing DHE staining followed by flow cytometry analysis of cells treated with all three compounds at 10 µM – a concentration that induced high levels of apoptosis in cancer cells and had minimal effects on non-cancer cells. Consistent with a role for mitochondrial-ROS in apoptosis, all three treatments induced ROS formation in cancer cells, but had minimal effects on non-cancer cells (**Fig. 1C**).

**Fig. 1.**
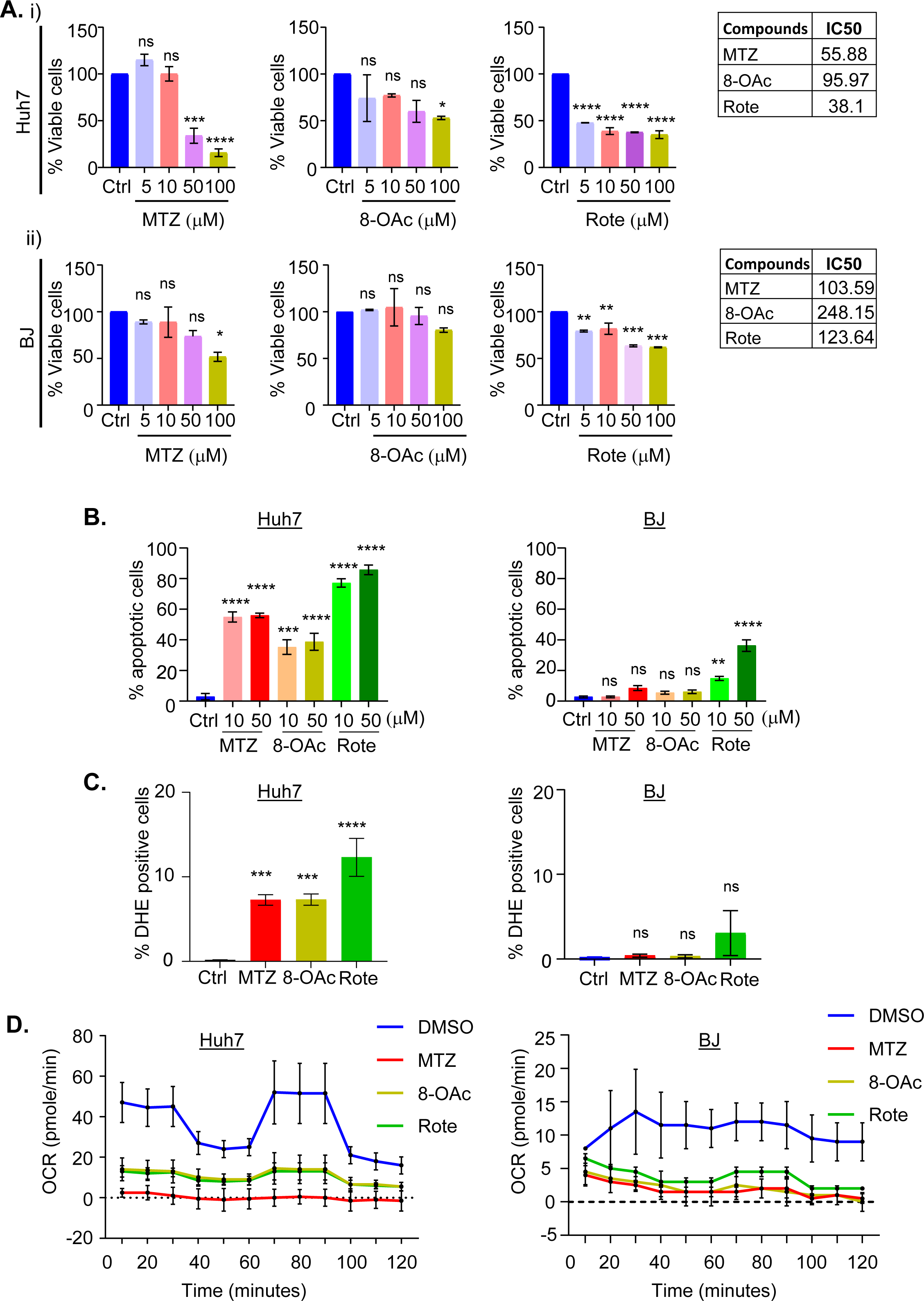
Cancer cells are more sensitive than non-cancer cells to mitochondrial complex I inhibitor MTZ and 8-OAc. (A) Cytotoxicity of (i) Huh-7 liver carcinoma cells and (ii) BJ fibroblast non-cancer cells by MTT assay after treatment with vehicle/DMSO or various concentrations of MTZ, 8-OAc, and Rote treatment for 24h. Corresponding chart shows the IC50 values of cells. (B) Bar graphs represent the percentage of early apoptotic cells analyzed by flow cytometry with annexin V and propidium iodide double staining after treatment with DMSO or indicated concentration of MTZ, 8-OAc, and Rote for 24h. (C) Graphs show the percentage of ROS-producing cells analyzed by flow cytometry with DHE staining after treating Huh-7 and BJ with DMSO or MTZ, 8-OAc, and Rote for 24h. (D) Graphs showing the inhibition of oxygen consumption rate after treating the indicated cells with 10μM of MTZ, 8-OAc, or Rote for 24h in comparison with vehicle control DMSO. Bar graphs represent mean and standard deviation. All statistical analysis was performed by one-way ANOVA using GraphPad Prism 10. ns = not significant, * = p < 0.03; ** = p <0.002; *** = p<0.0002; **** = p< 0.0001

In an effort to more thoroughly assess the impact of these compounds on general mitochondrial function, we measured mitochondrial respiration through monitoring oxygen consumption rate by Seahorse assay. The Seahorse assay results showed that all three compounds result in a significant decrease in mitochondrial respiration in both cancer and non-cancer cells (**Fig. 1D**). Interestingly, although all three compounds induce significant mitochondrial dysfunction in both cancer and non-cancer cells, we found that these compounds only induced mitochondrial fragmentation in cancer cells without affecting mitochondrial morphology in non-cancer cells (**Fig. s2B**). To further assess the general impact of these compounds on cellular health, we measured the impact of compound treatment on gene expression through RNA-seq analysis (**Fig. s3**). As expected, treatment of cancer cells with all three compounds resulted in significant gene expression changes in cell death, apoptosis, and cell cycle-related genes (**Tab. s1**). Perhaps most surprisingly, we failed to see any significantly differentially expressed genes when non-cancer cells are treated with the same concentration of these compounds. These data suggest that although all three compounds can robustly inhibit mitochondrial ETC function in both cancer and non-cancer cells, they induce dramatic transcriptional remodeling only in cancer cells, resulting in cancer-specific induction of cell death.

### Lower concentration MTZ and 8-OAc activate UPR^MT^ in worms but fail to extend lifespan

Next, to determine the impact of these compounds on mitochondrial function and organismal health in vivo, we moved into a *C. elegans* model system, particularly because UPR^MT^-mediated mitohormesis and impacts on longevity are well characterized in worms (Dutta, Garcia, and Higuchi-Sanabria 2022). Here, we exposed wild-type *C. elegans* to our compounds by supplementing nematode growth medium (NGM) with varying concentrations. We found that all three compounds do not impact development if animals are exposed to up to 3 µM concentrations from the 1^st^ larval stage (L1). However, concentrations above 3 µM resulted in a developmental delay, with MTZ and Rote having more profound effects than 8-OAc (**Fig. 2A**).

**Fig. 2.**
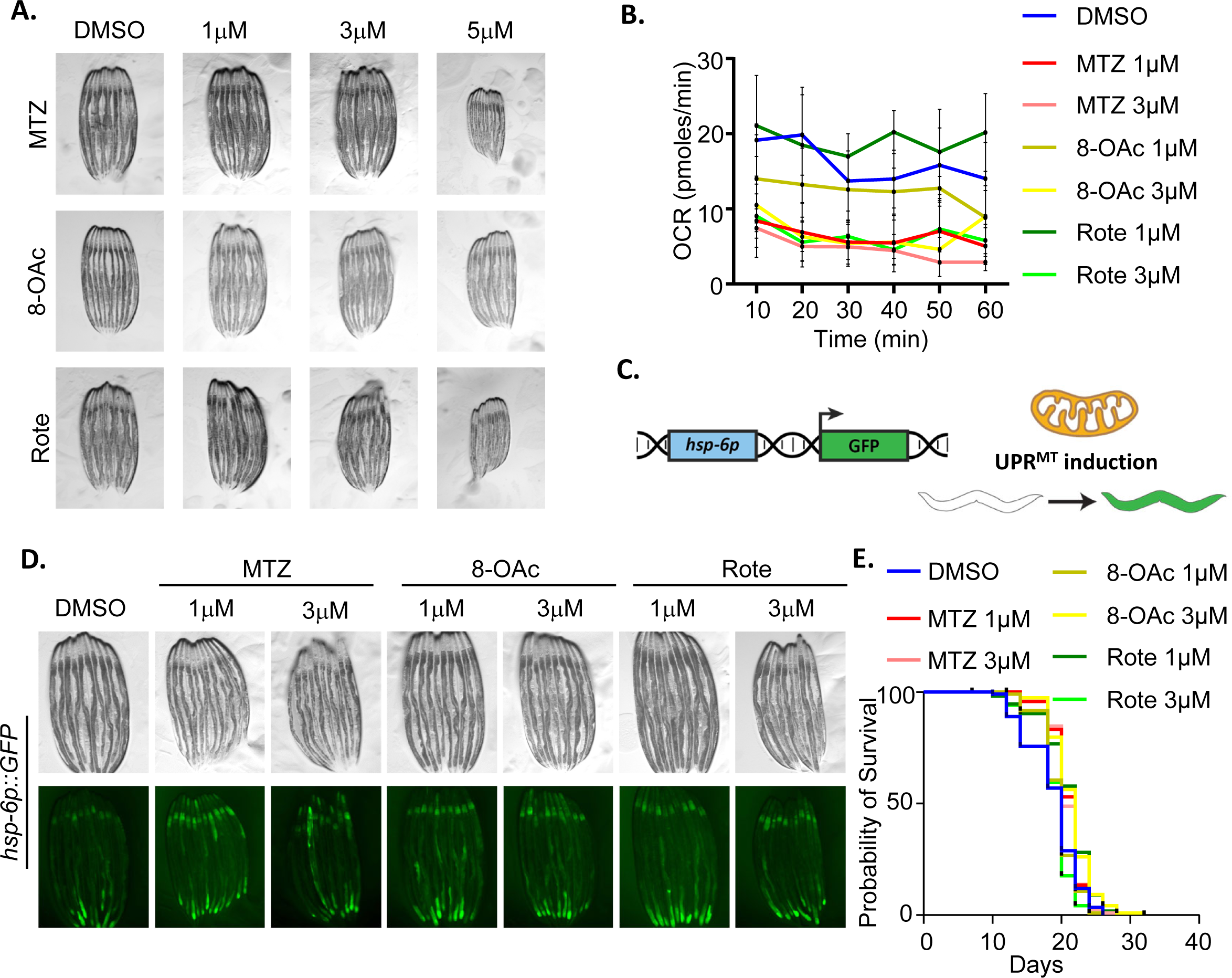
Low concentration of MTZ and 8-OAc activate UPR^MT^, but do not impact lifespan. (A) Representative images of wild-type N2 animals grown on DMSO controls or the indicated concentrations of MTZ, 8-OAc, or Rote from L1 for 3 days at 20 °C (i.e., day 1 adulthood for control conditions). (B) Seahorse analysis of mitochondrial respiration/OCR (pmol/min) of day 1 adult wild-type worms after growing them on indicated concentration of MTZ, 8-OAc or Rote from L1 until day 1 of adulthood. (C) Schematic representation of UPR^MT^ transcriptional reporter, *hsp-6p::GFP*. (D) Representative fluorescent micrograph of UPR^MT^ transcriptional reporter worms (*hsp-6p::GFP*) of day 1 adult wild-type animals grown on specified concentration of MTZ, 8-OAc, or Rote from L1. (E) Survival of wild-type N2 animals grown on DMSO control or indicated concentrations of MTZ, 8-OAc, or Rote from L1.

Since previous studies have shown that UPR^MT^-mediated mitohormesis generally involves early-life exposure to ETC inhibition during development (Dillin et al. 2002; Rea, Ventura, and Johnson 2007), we opted for 1 and 3 µM concentrations that allowed us to expose worms to these compounds throughout development. To first confirm that these compounds inhibited mitochondrial function in vivo, we again measured mitochondrial respiration rates using Seahorse assay. Interestingly, we found that although all three compounds significantly reduced oxygen consumption rates at 3 µM concentrations, only MTZ was sufficient to reduce respiration at the lower concentration of 1 µM (**Fig. 2B**). These data suggest that in *C. elegans*, MTZ has a more profound effect on mitochondrial respiration compared to 8-OAc and Rote.

Considering the established association of mitochondrial ETC inhibition and activation of the UPR^MT^, we next assessed the impact of MTZ and 8-OAc on the UPR^MT^ using the *hsp-6p::GFP* reporter (Yoneda et al. 2004) (**Fig. 2C**). We found that all three compounds induced UPR^MT^ activation at both the 1 and 3 µM concentrations (**Fig. 2D**). Importantly, this induction of UPR^MT^ required ATFS-1, the canonical transcription factor involved in UPR^MT^ activation (Nargund et al. 2012), as RNAi knockdown of *atfs-1* completely suppressed the UPR^MT^ induction upon compound treatment (**Fig. s4A**). We found that all three compounds also increased expression and nuclear localization of DVE-1::GFP, another robust reporter for UPR^MT^ activation (**Fig. s4B**) (Tian et al. 2016). Interestingly, all three compounds had profoundly different impacts on mitochondrial morphology in *C. elegans* (**Fig. s5**). Specifically, we found that all three compounds caused fragmentation of mitochondria in the muscle, but only MTZ caused fragmentation of mitochondria in the intestine at both concentrations. Rote only induced mitochondrial fragmentation in the intestine at the 3 µM concentration, whereas 8-OAc failed to induce mitochondrial fragmentation at either concentration. Finally, 8-OAc induced mitochondrial fragmentation in the hypodermis at both concentrations, whereas MTZ only induced mitochondrial fragmentation in the hypodermis at the 3 µM concentration and Rote failed to induce mitochondrial fragmentation at either concentration in the hypodermis. These data suggest that while all three compounds impact mitochondrial ETC function and induce the UPR^MT^, they display some differences in their downstream effects and may even have cell-type specific effects. Finally, to determine whether the inhibition of ETC and activation of UPR^MT^ was sufficient to drive mitohormesis and lifespan extension, we next measured lifespan of animals exposed to 1 and 3 µM concentration of the compounds from L1. Surprisingly, we did not observe any measurable increase in lifespan at either concentration for any of the compounds (**Fig. 2E**), suggesting that although UPR^MT^ is activated, it is not sufficient to drive mitohormesis.

### Higher concentration of MTZ and 8-OAc extend lifespan of adult worms

The absence of lifespan changes of worms at lower concentration of MTZ and 8-OAc led us to question whether higher concentrations of these compounds could potentially impact longevity. However, it is notable that we observed a developmental defect in worms when exposed to a concentration exceeding 3 μM beginning from the L1 stage (**Fig. 2A**). Therefore, to expose worms to higher concentrations of compounds, we opted to expose animals to plates containing 5 µM concentration of compounds at the beginning of day 1 of adulthood (**Fig. 3A**). Importantly, exposing animals to 5 µM concentration of these compounds at the day 1 adult stage was still sufficient to significantly reduce mitochondrial respiration rate (**Fig. 3B**). Exposure of animals to this higher concentration of compounds was sufficient to drive lifespan extension (**Fig. 3C**). To our surprise, we did not observe an induction of UPR^MT^ activity using the *hsp-6p::GFP* reporter (**Fig. 3D**) or DVE-1::GFP reporter (**Fig. s6A**). While these data may suggest that the lifespan extension is not through UPR^MT^-mediated mitohormesis, it is important to recognize that while the *hsp-6p::GFP* reporter is a robust method that allows for quickly testing for activation of UPR^MT^, it is a single-gene reporter and cannot entirely encompass the UPR^MT^ activation.

**Fig. 3.**
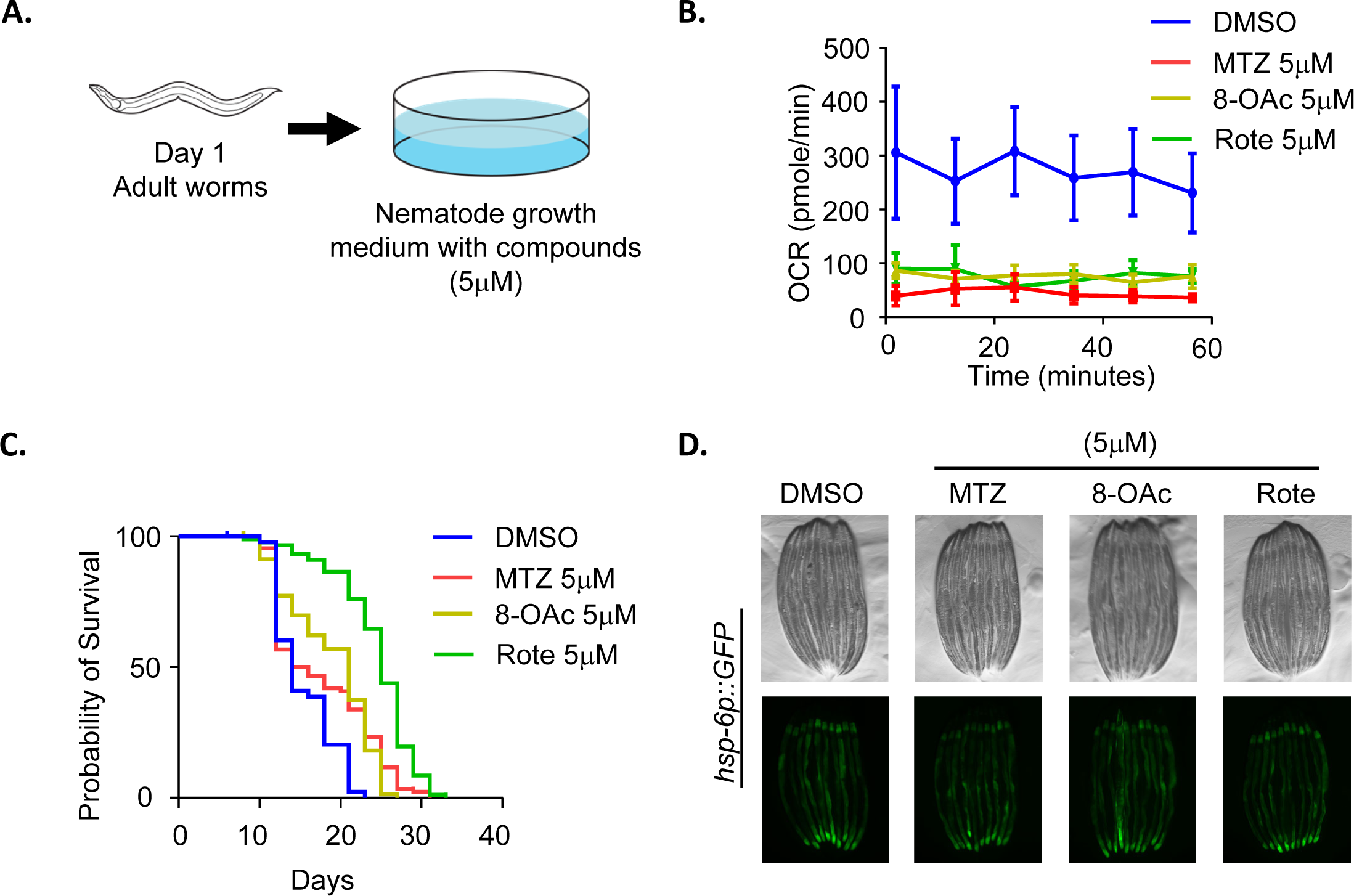
High concentrations of MTZ and 8-OAc promotes longevity. (A) Schematic of worms treated with high concentrations of compounds starting from day 1 of adulthood. (B) Seahorse analysis of mitochondrial respiration/OCR (pmol/min) of day 2 adult wild-type worms after growing them on 5 µM MTZ, 8-OAc, and Rote for 24 hours starting from day 1 of adulthood. (C) Representative fluorescent micrograph of UPR^MT^ transcriptional reporter worms (*hsp-6p::GFP*) of day 2 adult animals grown on 5 µM MTZ, 8-OAc, or Rote for 24 hours from day 1 of adulthood. (D) Survival of wild-type N2 animals grown on DMSO control or 5 µM MTZ, 8-OAc, or Rote from day 1 of adulthood.

Therefore, we performed a transcriptomic analysis by RNA-seq to more widely assess activation of the UPR^MT^. We found that MTZ treatment resulted in more widespread changes in transcriptome, while 8-OAc had the smallest effect (**Fig. 4A-B**). Importantly, genes related to UPR^MT^ showed significant changes for all three compounds, which confirms that they induce UPR^MT^, despite the lack of *hsp-6p::GFP* reporter induction (**Fig. 4C-D**). In addition, we did not see major changes in other stress response pathways including the unfolded protein response of the endoplasmic reticulum (UPR^ER^) or the heat-shock response (HSR) (**Fig. 3D**). These data were confirmed using reporters for the HSR (*hsp-16.2p::GFP*) and the UPR^ER^ (*hsp-4p::GFP*), both of which showed no induction upon treatment with MTZ, 8-OAc, or Rote (**Fig. s6B-C**). GO analysis of differentially expressed genes revealed many groups of genes that are consistent with a role for inhibition of mitochondrial function and mitohormesis pathways downstream of ETC dysfunction, including those involved in ETC function, response to reactive oxygen species, and mitochondrial transporter (**Fig. 4E**). Altogether, our transcriptomic analysis revealed that as expected, all three compounds induce UPR^MT^ pathways without affecting other stress response pathways.

**Fig. 4.**
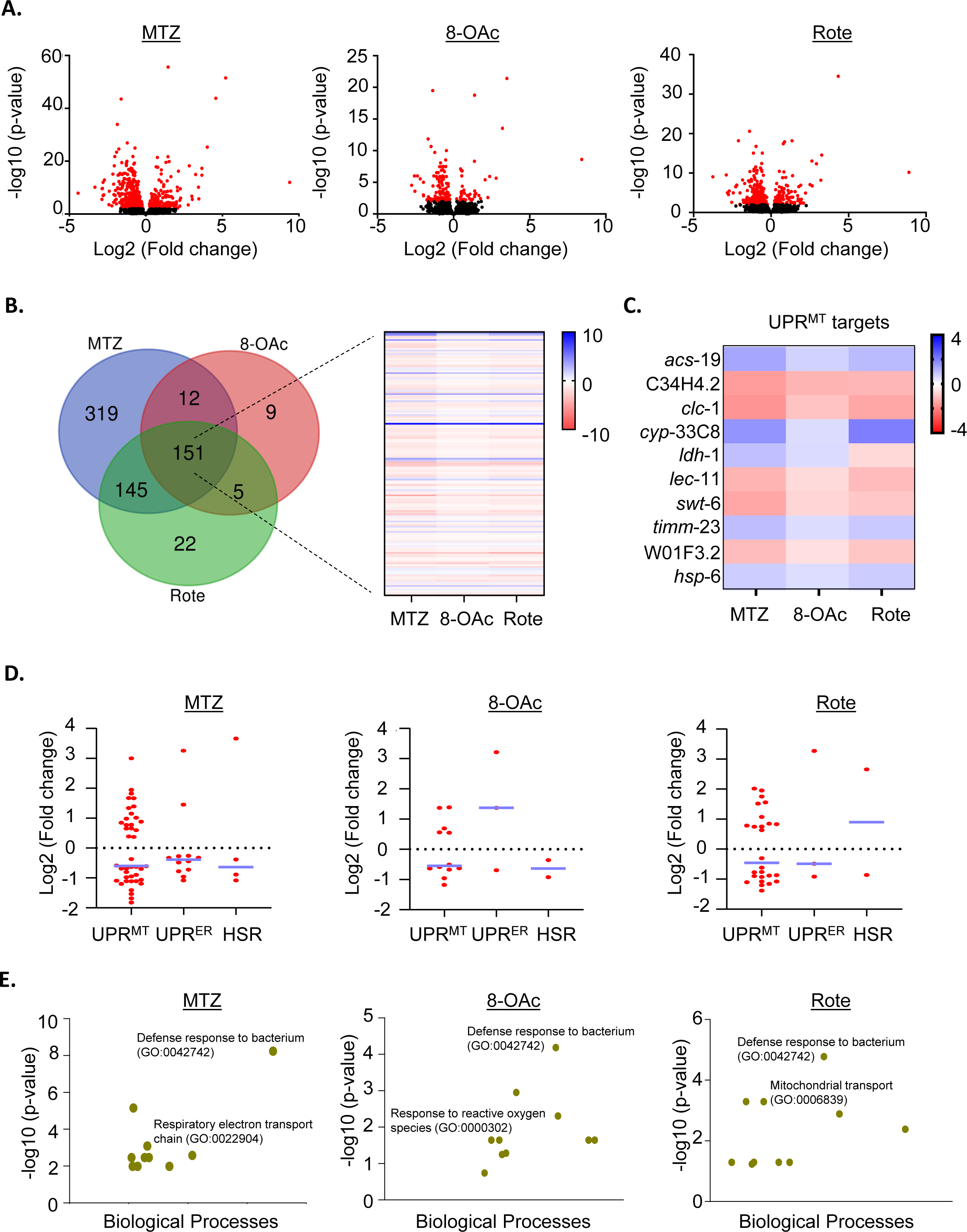
Transcriptomics analysis of MTZ, 8-OAc, and Rote. (A) Volcano plots representing the changes in gene expression in MTZ, 8-OAc, or Rote treated worms. For RNA-seq analysis, sterile *glp-4(bn2)* animals were grown at 22 °C until day 1 of adulthood and animals were moved onto compounds at day 1 and RNA was collected at day 2 of adulthood after 24 hours of growth on compounds. Data was analyzed on 3 biological replicates for each condition. Red dots indicate significantly differentially expressed genes, while black dots indicate genes that are not significant. (B) Venn diagram representing the overlap of significant differentially expressed genes in indicated treatment conditions. Heatmap represents the expression profile of common genes for indicated treatment conditions. (C) Heatmap showing the expression profile of UPR^MT^ target genes found common for MTZ, 8-OAc, or Rote treated conditions. (D) Representative graphs indicating the change in the gene expression of the indicated gene groups in MTZ, 8-OAc, or Rote treated condition as compared with DMSO control. Each dot represents a single gene and lines are median and interquartile range. (E) Gene enrichment analysis showing the most significant biological processes that are associated with the mentioned treatment conditions.

To directly test whether the lifespan extension observed after treatment with each compound is dependent on UPR^MT^ pathways, we performed lifespan experiments on animals with *atfs-1* knockdown where UPR^MT^ activation is suppressed. Interestingly, we found dramatic differences in the effect of all three compounds upon *atfs-1* knockdown. Specifically, MTZ and Rote treatment resulted in a decrease in lifespan with MTZ having more profound effects, suggesting that in the absence of a beneficial UPR^MT^ activation, not only do MTZ and Rote fail to extend lifespan, but they become toxic to worms. To our surprise, we found that the lifespan extension of 8-OAc is independent of *atfs-1* (**Fig. 5A**). Previous studies have revealed that the heat-shock transcription factor, HSF-1, is also required for activation of UPR^MT^ (Katiyar et al. 2020).

**Fig. 5.**
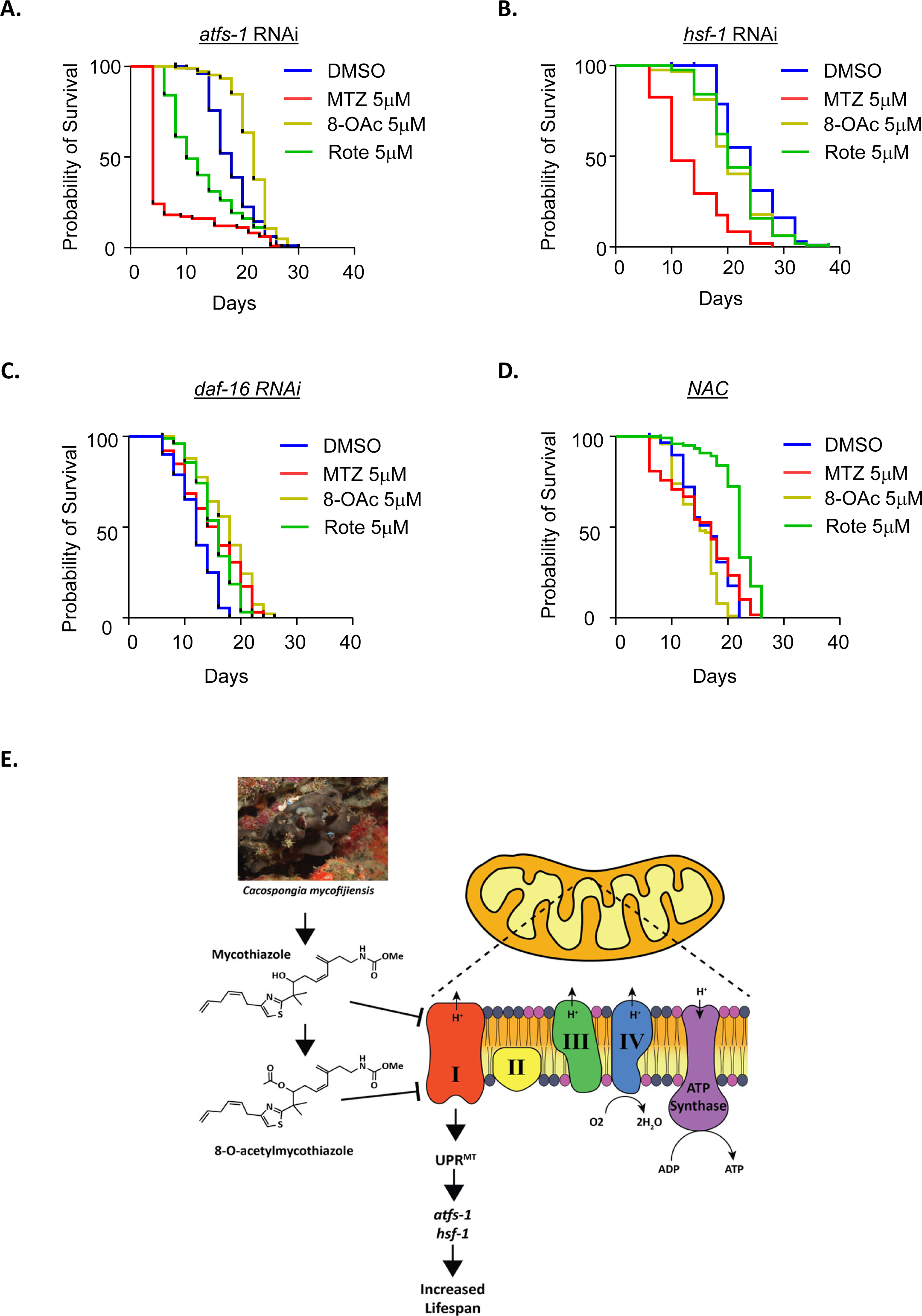
Lifespan extension of MTZ, 8-OAc, and Rote utilize different mechanism. (A) Survival of wild-type N2 animals grown on atfs-1 RNAi on DMSO control or 5 µM MTZ, 8-OAc, or Rote from day 1 of adulthood. (B) Survival of wild-type N2 animals grown on hsf-1 RNAi on DMSO control or 5 µM MTZ, 8-OAc, or Rote from day 1 of adulthood. (C) Survival of wild-type N2 animals grown on daf-16 RNAi on DMSO control or 5 µM MTZ, 8-OAc, or Rote from day 1 of adulthood. (A) Survival of wild-type N2 animals grown on NAC-supplemented plates containing DMSO control or 5 µM MTZ, 8-OAc, or Rote from day 1 of adulthood.

Therefore, we also tested the requirement of HSF-1 on lifespan extension and found that RNAi knockdown of *hsf-1* suppressed the lifespan extension found upon treatment by MTZ, 8-OAc, and Rote (**Fig. 5B**). Interestingly, similar to the loss of *atfs-1*, MTZ displayed a shortening of lifespan upon *hsf-1* knockdown, suggesting that in the context of *hsf-1* loss, MTZ is toxic.

Finally, since HSF-1 is also involved in oxidative stress response (Himanen et al. 2022) and our RNA-seq analysis revealed that 8-OAc resulted in changes in expression of antioxidant genes (**Fig. 4E**), we next sought to determine whether the lifespan extension found upon treatment with MTZ, 8-OAc, or Rote were dependent upon activation of antioxidant defense pathways. Therefore, we silenced *daf-16*, a crucial gene associated with the oxidative stress response pathway but did not observe any involvement of *daf-16* in the extension of lifespan under compound treatment (**Fig. 5C**). To further investigate the effect of redox homeostasis on longevity, we performed lifespan experiments after quenching reactive active species (ROS) within cells by using N-acetylcysteine (NAC) and observed that NAC suppressed the lifespan extension of both MTZ and 8-OAc treatment but did not affect the lifespan extension of Rote treatment. These data suggest that at least in the case of MTZ and 8-OAc, the synthesis of ROS, and likely the activation of downstream antioxidant pathways are involved in the longevity found upon treatment with these compounds.

## DISCUSSION

Gene level manipulation such as gene knockout or knockdown is a highly robust approach for studying the effect of a protein’s function in specific cellular events (Curtis and Nardulli 2009). However, these methods show several substantial limitations. First, animals with constitutive gene knockouts – especially for those knockouts that cause dramatic physiological changes – can pick up suppressors or adaptations, which can mask important phenotypes. Moreover, with certain strategies for knockdown or knockout, there are potential off-target effects, which can cause consequences from unintended genetic alternations (Höijer et al. 2022; Carroll 2019). Finally, the most prominent limitation may be in the challenge of performing genetic manipulations in highly complex systems, such as long-lived vertebrate systems or in human biology (Uddin, Rudin, and Sen 2020). In this context, small molecule chemical probe inhibitors, particularly those isolated from natural sources, offer distinct advantages over gene manipulation approaches (Weiss, Taylor, and Shokat 2007). They often offer high specificity in their mode of action, thus showing more reliability to target a particular protein or any specific cellular pathway. Importantly, they also provide temporal control for reversable modulation of a protein function. Finally, these chemical probes can be utilized in most model systems and in humans without the need for any genetic manipulations or interventions. As such, small molecule inhibitors have become a valuable source for therapeutic drug discovery (Beck et al. 2022). Modification in the structure of a known or previously characterized small molecule is also common practice in the field of drug discovery research to obtain new therapeutics with better pharmacological efficiency. Chemical modifications can improve the bioavailability of a small molecule through enhancing its permeability, absorption, distribution, and regulation of its excretion properties which in turn enhance the half-life of that molecule in the body. Alteration in the structure of a small molecule based on the knowledge of its target site can also increase the specificity of that molecule with its receptor, which can selectively reduce the off-target effects of that molecule. Modifications can also be done to provide more stability to the lead compound in diverse physiological conditions without hampering its basic properties (Y.-S. Ma et al. 2021; Xiong et al. 2021; Bech, Pedersen, and Jensen 2018).

In this study, we focused on compounds that target mitochondrial function. As an organelle involved in many important cellular processes including energy homeostasis, cellular metabolism, and programmed cell death, it is often a key target for the development of new therapeutics in drug development research. Mitochondrial function can be disrupted in various ways, including inhibition of different components of the ETC. Usually compounds that disrupt mitochondrial function result in toxic consequences like uncontrolled production of reactive oxygen species, which can induce cell death and lead to serious consequences including drug-induced liver injury and cardiotoxicity in humans (Barbier-Torres et al. 2017; Karamanlidis et al. 2013). Alternatively, there are several mitochondrial inhibitors such as metformin, nitric oxide, and arsenic trioxide, that are commonly studied in the clinic for potentially beneficial effects (Machado et al. 2023). Atovaquone, a specific inhibitor for ETC complex III also showed very promising anti-cancer activity in patients (Xiang et al. 2016). OPB-51602, a specific ETC complex I inhibitor displayed significant tumor regression in patients with secondary resistance to epidermal growth factor receptor inhibitors in a phase 1 clinical trial (Hirpara et al. 2019). The clinical utility of these inhibitors also extends beyond their anti-proliferative properties. Some potent ETC inhibitors such as the complex I inhibitor papaverine are used to reduce cellular oxygen consumption, which resulted in improved radiosensitivity to solid malignancies through tumor reoxygenation (Benej et al. 2018). However, it is noteworthy that the clinical trials of certain ETC inhibitors such as BAY87-2243, ASP4132, IACS-010759 were terminated due to their dose limiting toxicities (Xu et al. 2020; Janku et al. 2021; Yap et al. 2023). Thus, the study of additional mitochondrial inhibitors can be a valuable source for drug discovery.

Rotenone is another extensively studied molecule that inhibits the ETC, specifically complex I. It is mainly used as a pesticide in agriculture and aquaculture, but its use has raised major concern because of its off-target effects on other species in the agricultural field and aquatic ecosystem (Heinz et al. 2017). In human biology, it exhibits promising anticancer properties (Xiao et al. 2020), but in our study, we found that it showed significant toxicity to non-cancer cells. Previous studies have shown that MTZ and 8-OAc similarly exhibit significant activity against a wide range of cancer cell types (Johnson et al. 2020). In our study, we confirmed these findings and show that both MTZ and 8-OAc can induce ROS production, leading to cell death in liver cancer cells. Perhaps most important, we found that MTZ and 8-OAc exhibited negligible toxicity against non-cancer cell types at a concentration where Rote showed mild toxicity, suggesting this class has potential therapeutic utility based on its unique solid tumor selectivity.

We also found that all three compounds directly inhibited mitochondrial function in both cancer and non-cancer cells as measured by reduced oxygen consumption rate. However, these compounds only resulted in mitochondrial fragmentation in cancer cells and not non-cancer cells. Our data uncouples the effects of mitochondrial respiration on mitochondrial morphology and dynamics, at least in the case of cultured cells. It is important to acknowledge recent limitations observed in the field of mitochondrial research related to cell culture experiments. The rigid polystyrene substrate used in tissue culture plates can elevate mechanosignaling pathways in cells through activation of HSF1 (Tharp et al. 2021). Similarly, growth in high glucose conditions can also elevate HSF1 signaling and alter mitochondrial metabolism (Irshad et al. 2019). Thus, it is entirely possible that the effects we see in non-cancer cells may be obscured by the usage of both high glucose medium and growth on polystyrene, and further investigation is necessary to explore the specificity of these compounds on cancer cells in an in vivo system or organoid system that circumvents these issues.

Going forward, we opted to complement our cell culture system using the in vivo *C. elegans* model system. While *C. elegans* do not allow for cancer studies, they are an excellent model system for understanding the effects of mitochondrial inhibition on aging, particularly as a commonly used model for mitohormesis pathways (P. X. Chen et al. 2024). At first, we opted to use 1 and 3 µM concentrations of MTZ and 8-OAc due to developmental delays in worms exposed to higher concentrations of compounds. We opted for lower concentrations as numerous mitohormesis studies have shown the importance of exposure to mitochondrial stress during development, which would not have been possible at the concentrations that induced developmental defects. Shockingly, while we saw UPR^MT^ activation at these lower concentrations, there was no lifespan extension. Instead, we observed a significant lifespan extension when animals were exposed to these compounds at higher concentrations post-development. Several studies have shown that lifespan extension via mitohormesis through ETC inhibition require inhibition during development (Dillin et al. 2002; Rea, Ventura, and Johnson 2007). In addition, RNAi of various ETC complex subunits fail to extend lifespan in worms if applied in their adulthood. Notably, post-developmental inhibition of *atp-*3 can cause a remarkable induction in UPR^MT^ but shorten lifespan (Angeli et al. 2021). In contrast, our studies indicate that activation of a beneficial mitohormesis occurs after adulthood when ETC inhibition occurs via compounds. While it is entirely possible that this is due to different effects of compounds versus genetic intervention via RNAi, they highlight how mitohormesis via ETC inhibition are context dependent.

Interestingly, we did not observe activation of *hsp-6p::GFP* in animals exposed to high concentrations of MTZ, 8-OAc, or Rote, despite these animals exhibiting extended lifespans. These data complement studies that have shown that *hsp-6p::GFP* expression is not always correlated with lifespan extension in mitohormesis pathways (Bennett and Kaeberlein 2014). However, more comprehensive transcriptome analysis using RNA-seq did indeed show that exposure to all three compounds induced genes associated with canonical UPR^MT^, which highlight the fact that a single-gene reporter, while useful for preliminary analyses, cannot be used as final conclusive evidence for stress responses. Another interesting phenomenon we observed from our RNA-seq analysis was that 8-OAc displayed the least number of differentially expressed genes both in human cells and in *C. elegans* in comparison to MTZ and Rote. We hypothesize that this is potentially due to 8-OAc having a more specific effect on mitochondrial ETC function with less off-target effects. Indeed, genes differentially expressed under 8-OAc conditions are more related to inhibition of mitochondrial function including antioxidant defense pathways, while MTZ and Rote induced gene expression changes in a diverse array of seemingly unrelated pathways including defense response to bacterium, translational elongation process, sodium ion export from cell, cell morphogenesis, etc. (**Tab. s2**). It is possible that the increased stability of 8-OAc compared to MTZ can be partially responsible for these effects, as we have also observed that old (>6 mos.) – and likely partially degraded – MTZ (Johnson et al. 2020) can have toxic effects on *C. elegans* and cause premature death in animals. These findings highlight the importance of medicinal chemistry modifications to lead compounds for structural optimization to augment their chemical stability (shelf life) while maintaining their potency and selectivity required for specific modes of action.

Finally, one of our most interesting findings was the observation that MTZ, 8-OAc, and Rote utilized distinct downstream mechanisms to promote longevity in *C. elegans*. While the lifespan extension by MTZ was dependent on *atfs-1* and *hsf-1*, 8-OAc is only dependent on *hsf-1*. Numerous studies have implicated HSF-1 in mitochondrial stress response: William *et. al.* has shown in *C. elegans* that in response to ETC impairment, a mitochondria specific variant of HSF1 is activated by a specific dephosphorylation event, which results in the induction of specific heat shock proteins (Williams et al. 2020). Furthermore, another study in mammalian cells showed that HSF1 constitutively binds with the promoters of mitochondrial chaperones. In response to mitochondrial stress that occupancy remarkably enhanced and induces the production of mitochondrial chaperones (Katiyar et al. 2020). Finally, mitochondrial stress can prevent proteostasis collapse during aging by improving HSF-1 function at late age (Williams et al. 2020). Our observations align with these previous studies highlighting the overlap of mitohormesis and HSF-1, and its implications on aging. Still to be understood is whether the beneficial effects of HSF-1 in the context of MTZ and 8-OAc is through canonical UPR^MT^ pathways or some other function of HSF-1. Considering the lack of canonical HSR genes identified in our RNA-seq pathway, it is likely that the lifespan extension of MTZ is not through canonical HSR, but potentially through an ATFS-1/HSF-1 dependent activation of UPR^MT^. In the case of 8-OAc, it is possible that both HSF-1-mediated UPR^MT^ and HSF-1 driven antioxidant pathways may be involved in lifespan extension, as our RNA-seq revealed induction of several antioxidant pathways and NAC treatment suppressed lifespan extension in 8-OAc treated conditions. These findings underscore the complexity of UPR^MT^ regulation and advocate for further in-depth exploration of these pathways. Moreover, as a cautionary tale, our study highlights the critical importance of testing multiple compounds of the same chemotype as similar structures with seemingly identical effects may have some important differences on biological mechanisms of action.

In summary, this comprehensive study elucidates the efficacy of two highly potent and selective small molecule inhibitors of the ETC, MTZ and 8-OAc in both anti-aging and anti-cancer studies. These compounds hold promise as effective alternatives to existing small molecule inhibitor chemical probes, such as Rote, offering new insights in the domains of mitochondrial and aging-related research. The remarkable lack of off-target effects and higher stability of 8-OAc highlights the significance of how semisynthetic modifications to natural products can optimize the structure of lead compounds to provide improved stability, retained potency and selectivity. Ultimately the structure of 8OAc is poised to serve as a promising next generation chemical probe to accelerate the study of mitochondrial dynamics in chemical biology to potentially offer new avenues for the development of anti-cancer and anti-aging therapeutics.

## ACKNOWLEDGEMENTS

We thank Dr. Berenice Benayoun for assistance with computational analysis. We thank Dr. Sean Curran and members of the Curran lab, including Dr. Tripti Nair and Dr. Nicole Stuhr for technical assistance. M.A., A.H., S.A.J., and G.G. are supported by T32AG052374, T.C.T. and M.O. are supported by 1R25AG076400 from the National Institute on Aging, T.A.J. is supported by The Fletcher Jones Endowment fund of DUC, and R.H.S. is supported by R00AG065200 from the National Institute on Aging, Larry L. Hillblom Foundation Grant 2022-A-010-SUP, and the Glenn Foundation for Medical Research and AFAR Grant for Junior Faculty Award. M.V is supported by R01AG075130. Purify of compounds were assessed using the UCSC MS instrument supported by NSF CHE-1427922 to P.C. Some strains were provided by the CGC, which is funded by the NIH Office of Research Infrastructure Programs (P40 OD010440). Some gene analysis was performed using Wormbase, which is funded on a U41 grant HG002223.

## AUTHOR CONTRIBUTIONS

N.D., G.G., T.A.J., and R.H.S. designed and oversaw all experiments. N.D. performed or oversaw all biological experiments (i.e., *C. elegans* and human cell culture) and prepared the manuscript and figures. J.A.G., S.F.O., J.D.M., J.G.C., M.C.C, M.N.R., H.S., and T.A.J. designed and performed all chemistry experiments including purification and quality control of all compounds. P.C. supplied repository crude extracts of *C. mycofijiensis*. T.C.T. and A.A. prepared all drug-containing plates for *C. elegans* workflow and assisted with *C. elegans* experiments. A.H., S.J.S., and M.A.T. assisted with all cell culture experiments. M.A., S.H., M.O., A.A., M.Vega, and R.H.S. assisted with lifespan experiments. N.D. and G.G. performed all computational analysis. M.Vermulst. provided expertise for cell culture experiments. All authors edited and approved the manuscript.

## DATA AVAILABILITY

All data required to evaluate the conclusions in this manuscript are available within the manuscript and Supporting Information. All strains synthesized in this manuscript are derivatives of N2 or other strains from CGC and are either made available on CGC or available upon request. All raw RNA-seq datasets are available through Annotare 2.0 Array Express Accession E-MTAB-13546.

