## Supplemental Information for "Investigating impacts of marine sponge derived mycothiazole and its acetylated derivative on mitochondrial function and aging"

### Supplemental Figures and Figure legends

**A.**

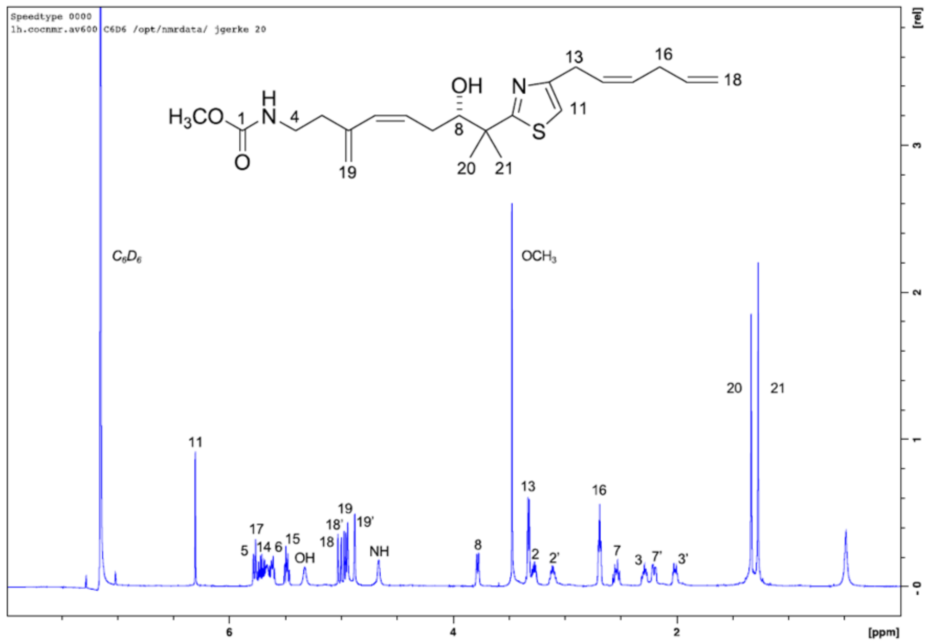


**B.**

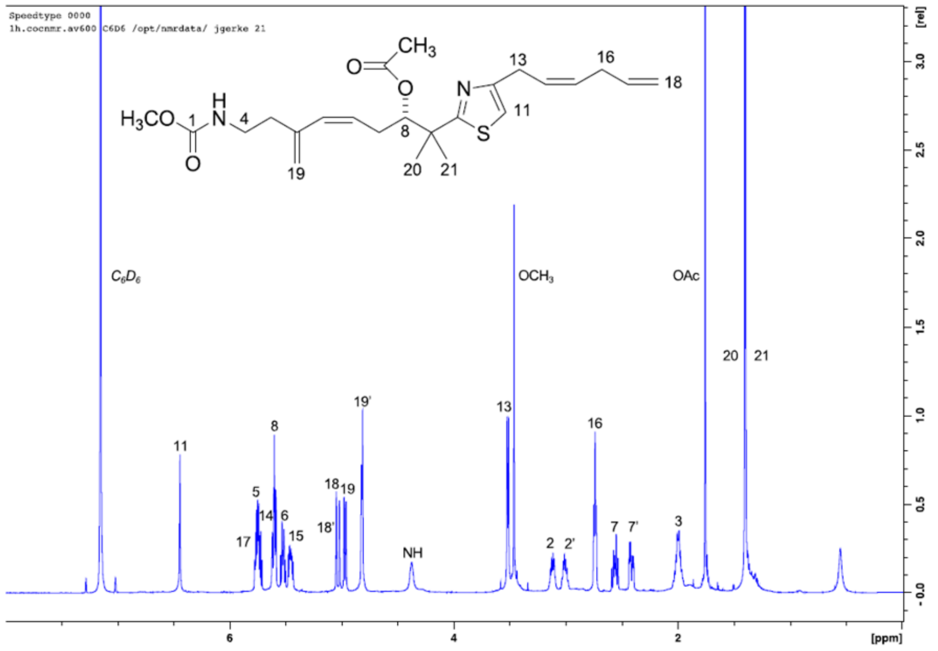


**Fig.s1. NMR analysis of MTZ and 8-OAc.**

A.

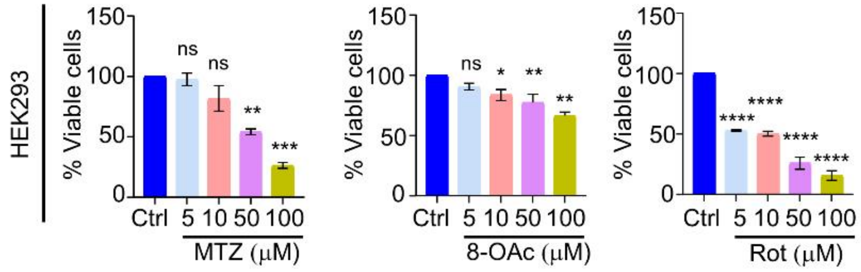


| Compounds | IC50 |
|---|---|
| MTZ | 62.95 |
| 8-OAc | 155.61 |
| Rote | 30.97 |

B.

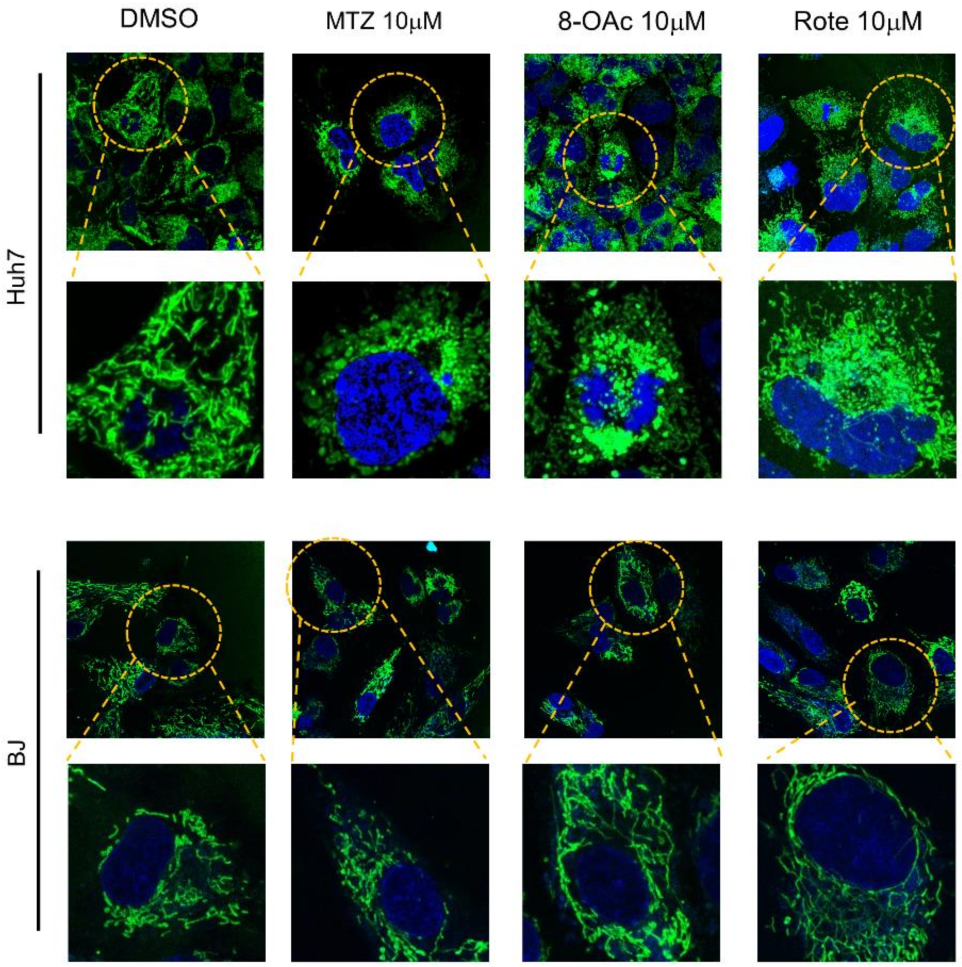


**Fig. s2. MTZ, 8-OAc, and Rote preferentially target cancer cells and not non-cancer cells.** (A) Cytotoxicity of HEK293T non-cancer cells by MTT assay after treatment with vehicle/DMSO or various concentrations of MTZ, 8-OAc, and Rote treatment for 24h. Corresponding chart shows the IC50 values of cells. (B) Representative images of Huh7 (top) and BJ fibroblast (bottom) cells treated with DMSO or 10 µM MTZ, 8OAc, and Rote treated with DAPI and mitotracker green as described in Materials and Methods.

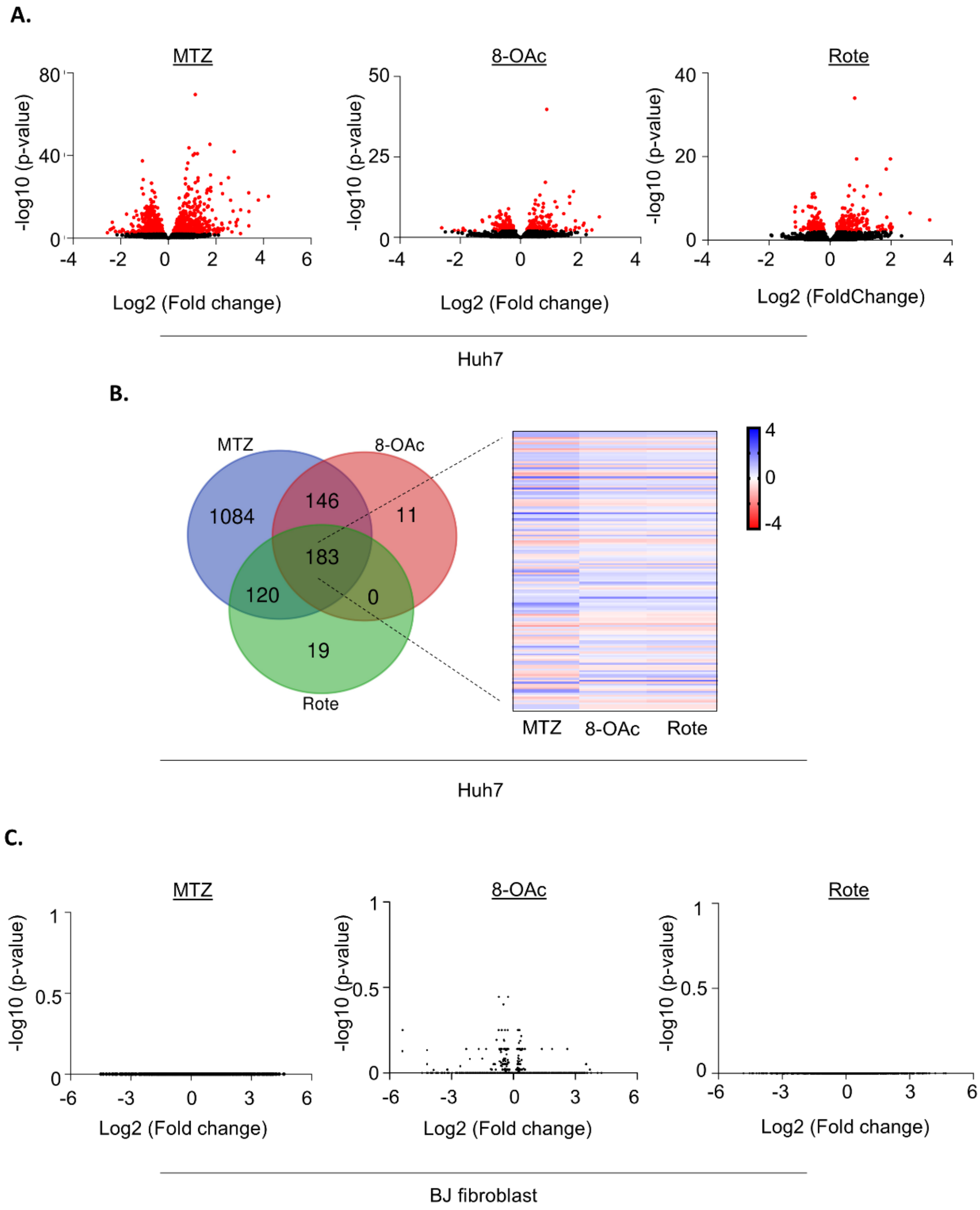


**Fig. s3. MTZ, 8-OAc, and Rote alter gene expression in cancer cells, but not in non-cancer cells.** (A) Volcano plots representing changes in gene expression in Huh7 liver carcinoma cells treated with 10 µM MTZ, 8-OAc, and Rote for 24 hours. Red dots indicate significantly differentially expressed genes. Data was analyzed on 3 biological replicates per condition. (B) Venn diagram representing the overlap of significantly differentially expressed genes in indicated treatment conditions. (C) Heatmap showing the expression of overlapping differentially expressed genes shared between MTZ, 8-OAc, and Rote. (D) Volcano plots representing changes in gene expression in BJ fibroblasts treated with 10 µM MTZ, 8-OAc, and Rote for 24 hours.

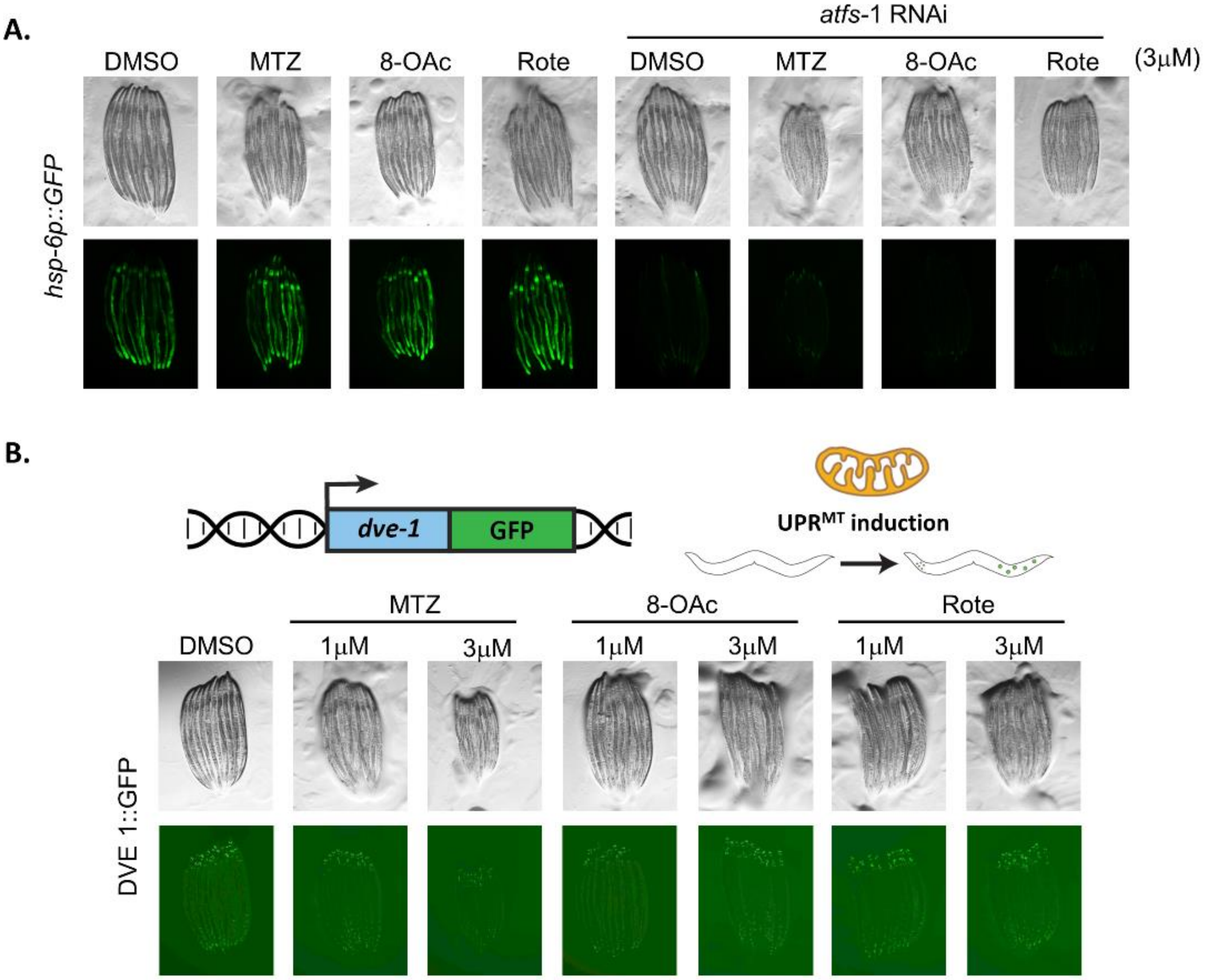


**Fig. s4. Low concentrations of MTZ, 8-OAc, and Rote induce UPR$^{MT}$ in an *atfs-1* dependent manner.** (A) Representative fluorescent micrograph of UPR$^{MT}$ transcriptional reporter worms (*hsp-6p::GFP*) of day 1 adult animals grown on empty vector or *atfs-1* RNAi plates supplemented with 3 µM MTZ, 8-OAc, or Rote for 24 hours from L1. (B) Representative fluorescent micrographs of day 1 adult DVE-1::GFP worms grown on NGM plates supplemented with DMSO or indicated concentrations of MTZ, 8-OAc, or Rote for 24 hours from L1.

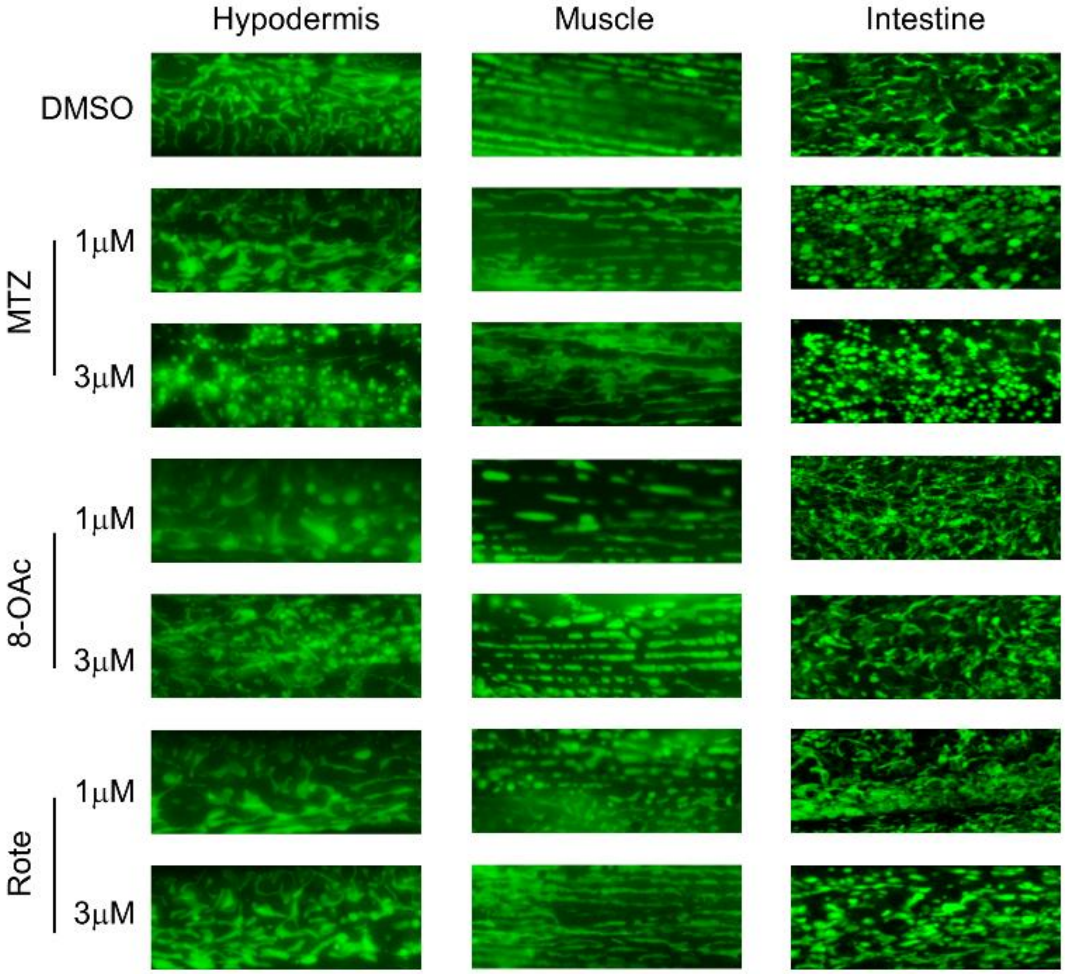


**Fig. s5. Low concentrations of MTZ, 8-OAc, and Rote induce mitochondrial fragmentation.** Representative fluorescence micrographs of MLS::GFP expressed in the hypodermis (*col-19p*), muscle (*myo-3p*), and intestine (*gly-19p*) of day 1 adult worms grown on NGM plates supplemented with DMSO or indicated concentrations of MTZ, 8-OAc, or Rote for 24 hours from L1.

**A.**

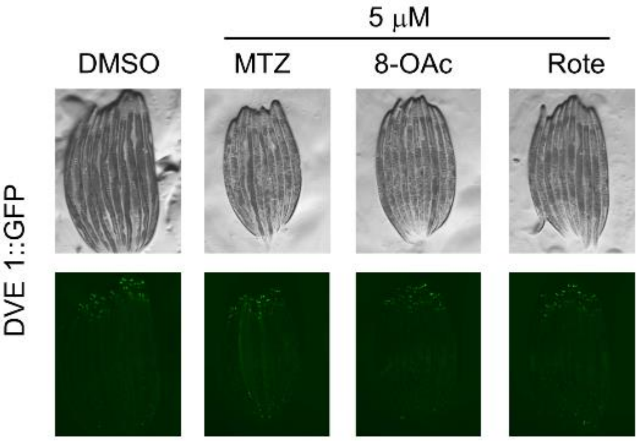
5 μM
DMSO
MTZ
8-OAc
Rote
DVE 1::GFP

**B.**

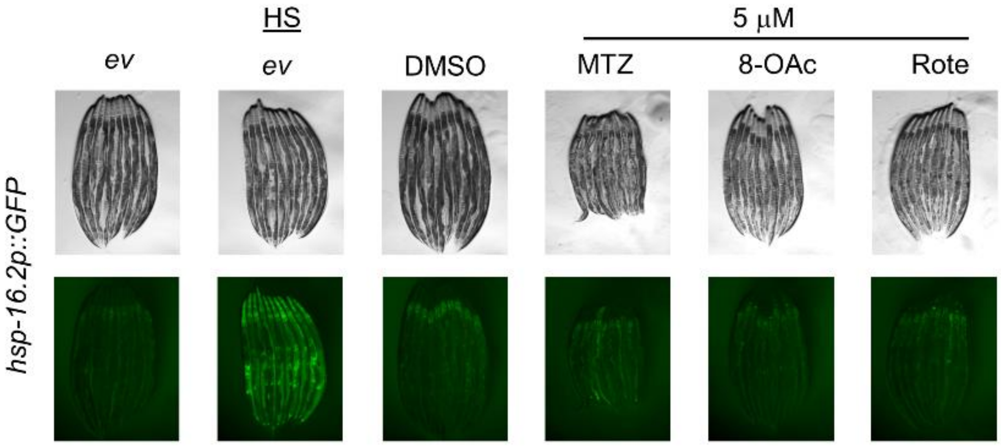
HS
5 μM
*ev*
*ev*
DMSO
MTZ
8-OAc
Rote
*hsp-16.2p::GFP*

**C.**

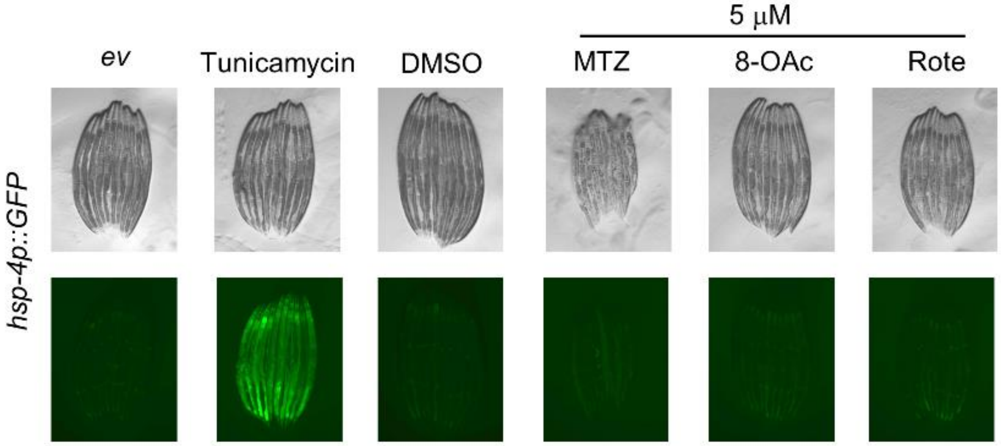
5 μM
*ev*
Tunicamycin
DMSO
MTZ
8-OAc
Rote
*hsp-4p::GFP*

**Fig. s6. High concentrations of MTZ, 8-OAc, and Rote do not induce reporters for stress responses.** (A) Representative fluorescent micrographs of DVE-1::GFP of day 2 adult animals grown on 5 µM MTZ, 8-OAc, or Rote for 24 hours from day 1 of adulthood. (B) Representative fluorescent micrograph of HSR transcriptional reporter worms (*hsp-16.2p::GFP*) of day 2 adult animals grown on 5 µM MTZ, 8-OAc, or Rote for 24 hours from day 1 of adulthood. Heat-shock for 2 hours at 34°C is used as a positive control. (C) Representative fluorescent micrograph of $UPR^{ER}$ transcriptional reporter worms (*hsp-4p::GFP*) of day 2 adult animals grown on 5 µM MTZ, 8-OAc, or Rote for 24 hours from day 1 of adulthood. Growth on tunicamycin plates is used as a positive control.
